# Bystander CD4^+^ T cells infiltrate human tumors and are phenotypically distinct

**DOI:** 10.1101/2020.07.15.204172

**Authors:** Yannick Simoni, Shamin Li, Summer Zhuang, Antja Heit, Si-Lin Koo, I-Ting Chow, William W. Kwok, Iain Beehuat Tan, Daniel S.W. Tan, Evan W. Newell

## Abstract

Tumor-specific T cells likely underpin effective immune checkpoint-blockade therapies. Yet, most studies focus on Treg cells and CD8^+^ tumor-infiltrating lymphocytes (TILs). Here we study CD4^+^ TILs in human lung and colorectal cancers and observe that non-Treg CD4^+^ TILs average more than 70% of total CD4^+^ TILs in both cancer types. Leveraging high dimensional analyses including mass cytometry and single-cell sequencing, we reveal that CD4^+^ TILs are heterogeneous at both gene and protein levels, within each tumor and across patients. Consistently, we find different subsets of CD4^+^ TILs showing characteristics of effectors, tissue resident memory (Trm) or exhausted cells (expressing PD-1, CTLA-4 and CD39). In both cancer types, the frequencies of CD39^−^ non-Treg CD4^+^ TILs strongly correlate with frequencies of CD39^−^ CD8^+^ TILs, which we and others have previously shown to be enriched for cells specific for cancer-unrelated antigens (bystanders). *Ex-vivo*, we demonstrate that CD39^−^ CD4^+^ TILs can be specific for cancer unrelated antigens, such as HCMV epitopes. Overall, our findings highlight that CD4^+^ TILs cells are not necessarily tumor-specific and suggest measuring CD39 expression as a straightforward way to quantify or isolate bystander CD4^+^ T cells.

**Graphical abstract:** 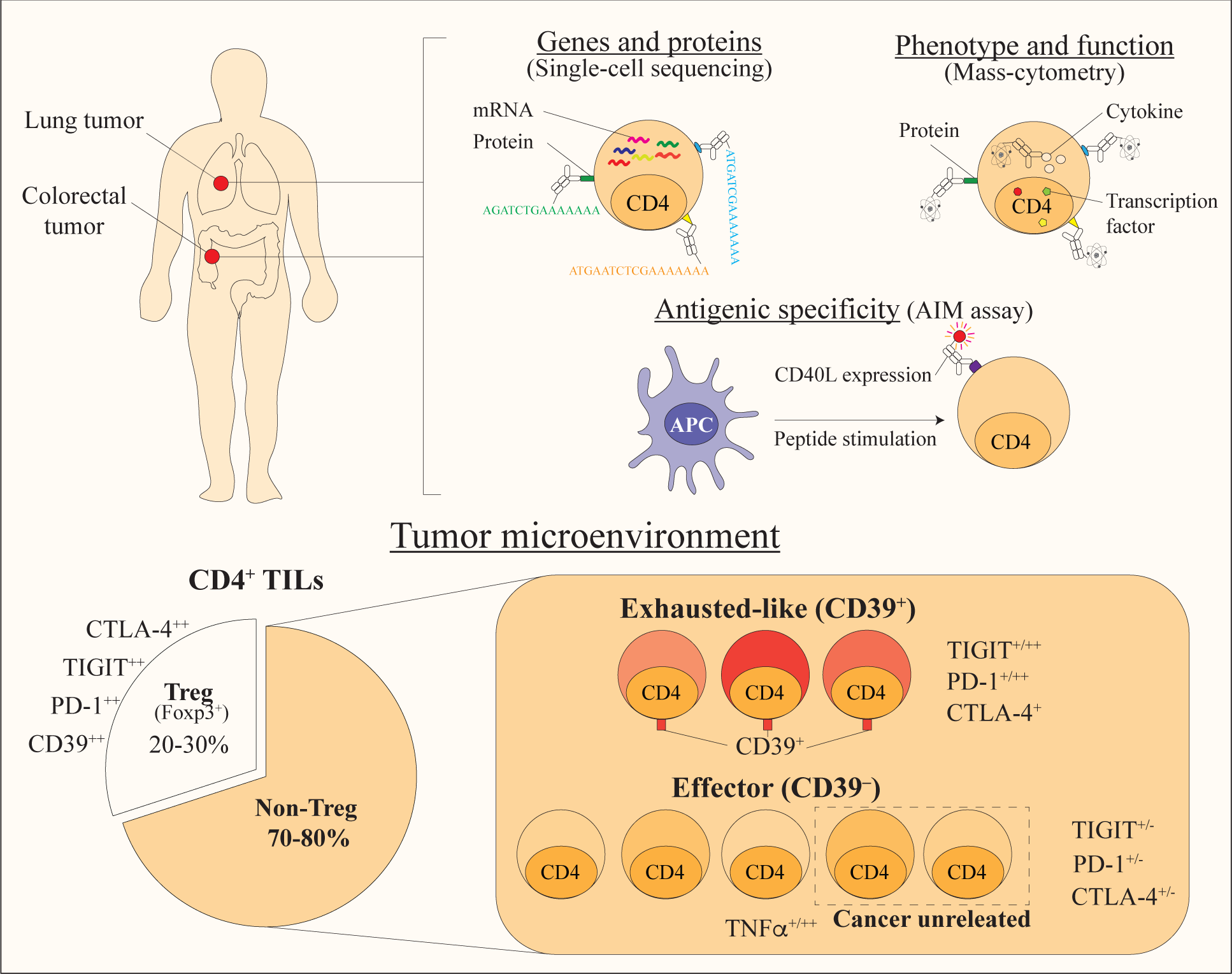

## Introduction

Numerous studies have established the importance of T cells in controlling cancer (1). Nonetheless, tumors can escape this immune surveillance by diverse mechanisms (2). As various forms of cancer therapy exist, immunotherapy is rapidly evolving and is proved to be remarkably effective at restoring T cell mediated immune responses. Strategies include immune checkpoint blocking receptors (i.e anti-CTLA4 or anti-PD1 (3), autologous T cell transfer (4), as well as therapeutic cancer vaccines (5). However, the efficacy of these therapies is unpredictable and only some patients respond well to the treatments (6). Therefore, a better understanding of T cell biology – CD8 and CD4 – in the tumor microenvironment is urged to improve cancer therapies. Recently, we showed in the context of human colorectal and lung cancers that CD8^+^ tumor-infiltrating lymphocytes (TILs) are not only specific for tumor antigens but can also recognize a wide range of cancer-unrelated epitopes (called bystander CD8^+^ TILs) (7). We suggested that measuring CD39 expression could be a straightforward way to quantify or isolate bystander CD8^+^ T cells and could be a potential biomarker for immunotherapy (7). These observations have been confirmed in different cancer types (8-11).

Although CD4^+^ TILs are also involved in tumor responses, most studies have focused on the role of FoxP3-expressing regulatory T cells (Treg) in cancer (12-14). Treg cells suppress tumor immunity by various mechanisms including: 1) Disruption of the metabolic pathway (i.e. CD39 expression), 2) Modulation of dendritic cells function (i.e. CTLA-4 expression), 3) Production of anti-inflammatory molecules (i.e. IL-10, TGFβ), 4) Induction of apoptosis (15). Abundant Treg infiltration into tumors is strongly associated with poor prognosis in multiple cancer types (13, 16). Because of their deleterious role, several molecules have been developed to target specifically these cells in human cancer (e.g. anti-CTLA-4, anti-CD25) (17-21).

Importantly, a large proportion of CD4^+^ TILs are made up of non-Treg cells. Studies in mice have shown that these cells play a key role in anti-tumor responses (22). By producing IFNγ, they induce an up-regulation of MHC class I and II expression by tumor cells and dendritic cells (DC) (23). Production of IFNγ by CD4^+^ TILs also induce expression of chemokines supporting homing of CD8^+^ T cells to the tumor site (e.g. CXCL10) (23). Activated CD4^+^ T cells express CD40L by which they can activate DC, and support CD8^+^ T cells priming and memory formation (23). They can have a cytotoxic function and directly kill tumor cells as well (24). Based on these observations, developing CD4-based therapeutic vaccination and/or adoptive cell therapies by targeting tumor-specific CD4^+^ T cells would be essential (22, 25-28). The limited number of tools that allow studying non-Treg CD4^+^ TILs (i.e. MHC class II tetramers, *in-vitro* assays) had so far made this population poorly characterized, compared to CD8^+^ TILs and Treg cells. Uncovering the role of these cells in the tumor microenvironment would thus help design new strategies to manipulate them and improve immunotherapy efficiency. Here we study CD4^+^ TILs in human colorectal cancer (CRC) and non-small cell lung cancer (NSCLC) using complementary high-dimensional single-cell analysis (single-cell sequencing, mass-cytometry) and *in-vitro* stimulation assay. Our findings highlight that non-Treg CD4^+^ TILs are heterogeneous and can be specific for cancer unrelated antigens, just as observed for CD8^+^ TILs, and these cells lack expression of CD39. Taken together, we hypothesize that CD39 expression is a straightforward way to quantify or isolate bystander CD4^+^ TILs, thus opening new diagnostic and therapeutic avenues.

## Results

### Single-cell Protein/mRNA sequencing reveals the heterogeneity of CD4^+^ TILs

In order to comprehensively examine CD4^+^ tumor infiltrating T cells (TILs), we leveraged the use of a recent single-cell sequencing technology that allows simultaneous analysis of surface protein and mRNA expression at the single-cell level (29, 30) (Figure 1A). The surface protein antibodies panel for instance included a broad range of markers associated with T cell differentiation, activation, tissue residency, and dysfunction/exhaustion status (co-stimulatory and co-inhibitory receptors). Prior to the single-cell experiment, tumor cells (Epcam^+^), myeloid cells (CD14^+^) and B cells (CD19^+^) were depleted (See Methods). To assess the composition of the total sequenced cells, we performed a Uniform Manifold Approximation and Projection (UMAP) based on surface protein expressions (31). UMAP is a dimension reduction algorithm that performs a pair-wise comparison of the cellular phenotypes to optimally plot similar cells close to each other (31). For our analysis, 48 surface parameters, or dimensions, were reduced into two dimensions (UMAP1 and UMAP2). This visualization allowed us to easily identify a population of contaminating tumor cells (CD45^−^), NK cells (CD3^−^), CD8^+^ TILs (CD3^+^ CD8^+^) and our cells of interest: CD4^+^ TILs (CD3^+^ CD4^+^) (Figure 1B, 1C and S1A). The phenotypic profile that we observed for these cells was depicted in a heatmap showing expression intensities of surface markers (Figure 1D). As for the CD4^+^ TILs, we observed a first subset characterized by markers associated with Treg cells (CD25^+^ CD39^+^ ICOS^+^ GITR^+^). Interestingly, the remaining CD4^+^ TILs could be divided based on their expression of CD39, a marker associated with chronic TCR stimulation (32)(Figure 1D and S1B). Based on their phenotypic difference, we studied each population at the transcriptomic level. As expected, the CD4^+^ subset defined phenotypically as Treg cells expressed their signature genes (i.e. *FOXP3, CTLA4, DUSP4*) (33). Interestingly, both Treg and CD39^+^ CD4^+^ TILs expressed *IL32*, a cytokine which enhances NK cell sensitivity and cytotoxicity against tumor cells (34). Furthermore, compared to CD39^+^, CD39^−^ CD4^+^ TILs expressed more of *TNF* transcript, suggesting a non-exhausted profile. Our results did not show significant differences in *IFNG*, cytotoxicity or chemokine expression between the different subsets of CD4^+^ TILs (Figure 1E).

**Figure 1.**
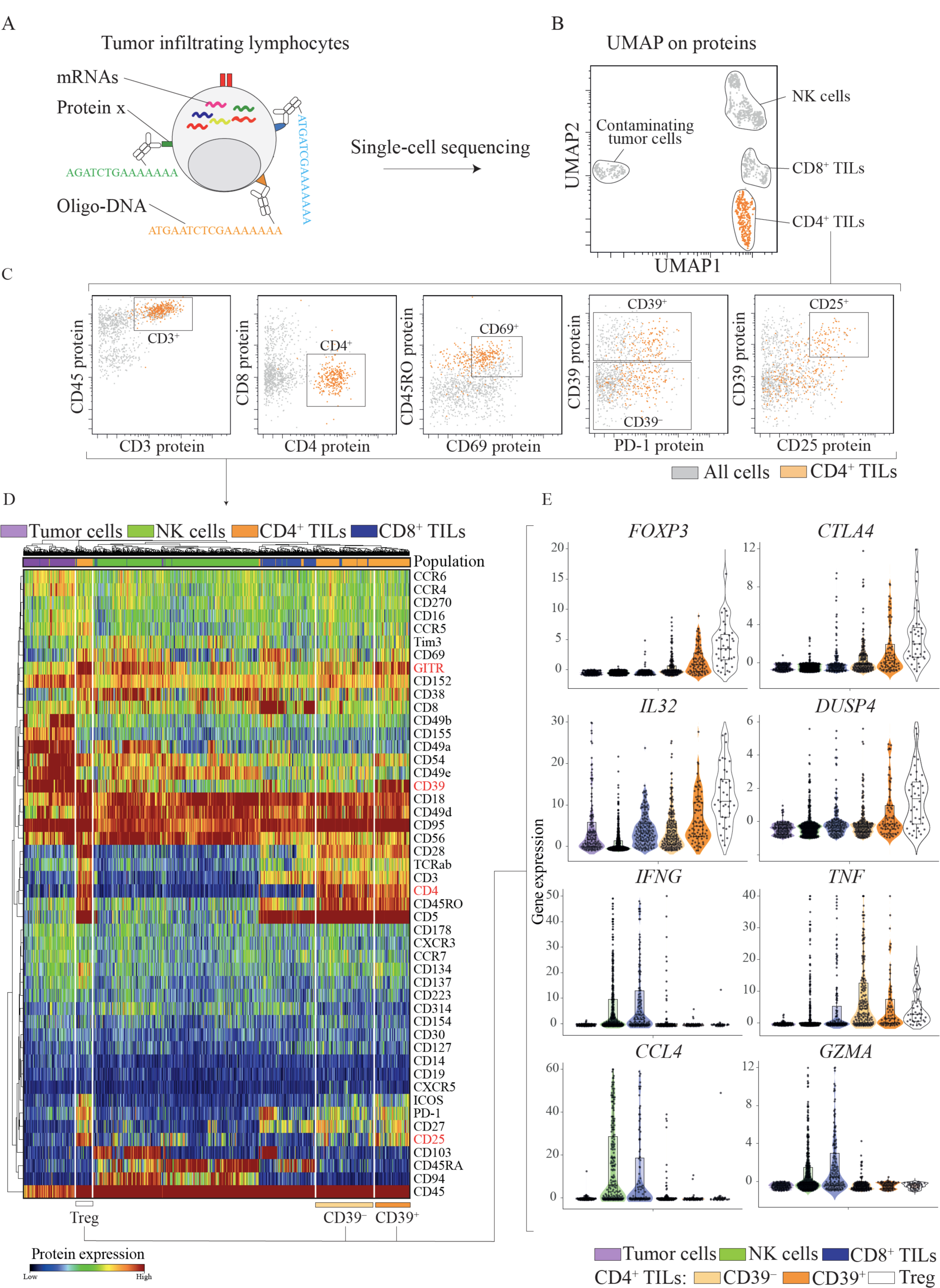
Phenotypic and transcriptomic heterogeneity of CD4+ TILs. **A**. Schematic for targeted mRNA sequencing combined with BD AbSeq Rhapsody system that allows analysis of both genes and surface proteins at the single cell level. **B**. UMAP plot on total cells sequenced from a colorectal tumor sample (patient 1211), after depletion for tumor cells (EpCAM+), B cells (CD19+) and myeloid cells (CD33+) before the experiment. **C**. Single-cell sequencing data showing expression of selected oligo-tagged surface protein markers for CD4+ TILs (orange) and total cells (grey). **D**. Heatmap from scRNAscq data depicting protein expression of all sequenced cells from this tumor at the single-cell level. Each cell population is identified by a different color code as indicated. **E**. Violin plots comparing expression of selected genes in different cell subsets of interests.

Taken together, our results indicated that CD4^+^ TILs were composed of heterogeneous populations that could be divided into Treg, CD39^+^ and CD39^−^ non-Treg CD4^+^ T cells. Additional samples will be needed to validate these observations in other patients.

### CD4^+^ TILs are composed of a majority of non-Treg cells with a contrasted phenotypic profile

We next investigated whether the heterogeneity we observed were consistent across patients and different tumor types. For that purpose, we profiled a cohort of patients with Non-small cell lung cancer (NSCLC, n=28) and colorectal cancer (CRC, n=51). We developed a mass cytometry panel consisting of 38 heavy metal-labelled antibodies to identify and characterize CD4^+^ TILs with markers of tissue residency, activation and inhibitory receptors (Table S1). We distinguished Treg and non-Treg cells based on the expression of FoxP3 (Figure 2A). With only 35% and 24% of CD8^+^ TILs in NSCLC and CRC respectively, the majority of CD3^+^ TILs were composed of CD4^+^ TILs (Figure 2B and 2C). Non-Treg CD4^+^ TILs accounted for a higher proportion of the CD4^+^ TILs as compared to Treg CD4^+^ TILs, with a mean frequency of 78.8% vs. 19% in NSCLC and 66% vs. 35% in CRC, supporting the importance of studying this population in tumor immune response (Figure 2B and C). All non-Treg CD4^+^ TILs displayed a memory or effector phenotype (CD45RO^+^ – 95.7%) and many expressed the activation/tissue residency marker CD69 (CD69^+^ – 77%), excluding a blood contamination for most of these cells (Figure 2D and S2A). Expression of activation markers and inhibitory receptors varied greatly in these cohorts, indicating an important phenotypic diversity of CD4^+^ TILs between patients (Figure 2D and S2A). Non-Treg CD4^+^ TILs expressed co-stimulatory receptors, such as CD28, CD38, ICOS but only a small fraction expressed CD127 (17.1%). Interestingly, some non-Treg CD4^+^ TILs expressed CD25 (26.7%), suggesting that the use of CD25 and CD127 alone to identify Treg cells in the context of tumor infiltrates could lead to a contamination by non-Treg CD4^+^ TILs (i.e. Foxp3^−^) (Figure 2D, S2A and 2E). More interestingly, non-Treg CD4^+^ TIL cells also expressed hallmarks of “exhausted” cells at different levels between patients. Expression of inhibitory receptors associated with chronic antigen stimulation such as TIGIT (56.9%), PD-1 (71.6%), CTLA-4 (29.6%) suggested a role for these cells in tumor immunity (Figure 2D and S2A). Of note, frequency of CD39^+^ non-Treg CD4^+^ TILs (38.2%) was very heterogeneous, ranging from 4.6% to 70%.

**Figure 2.**
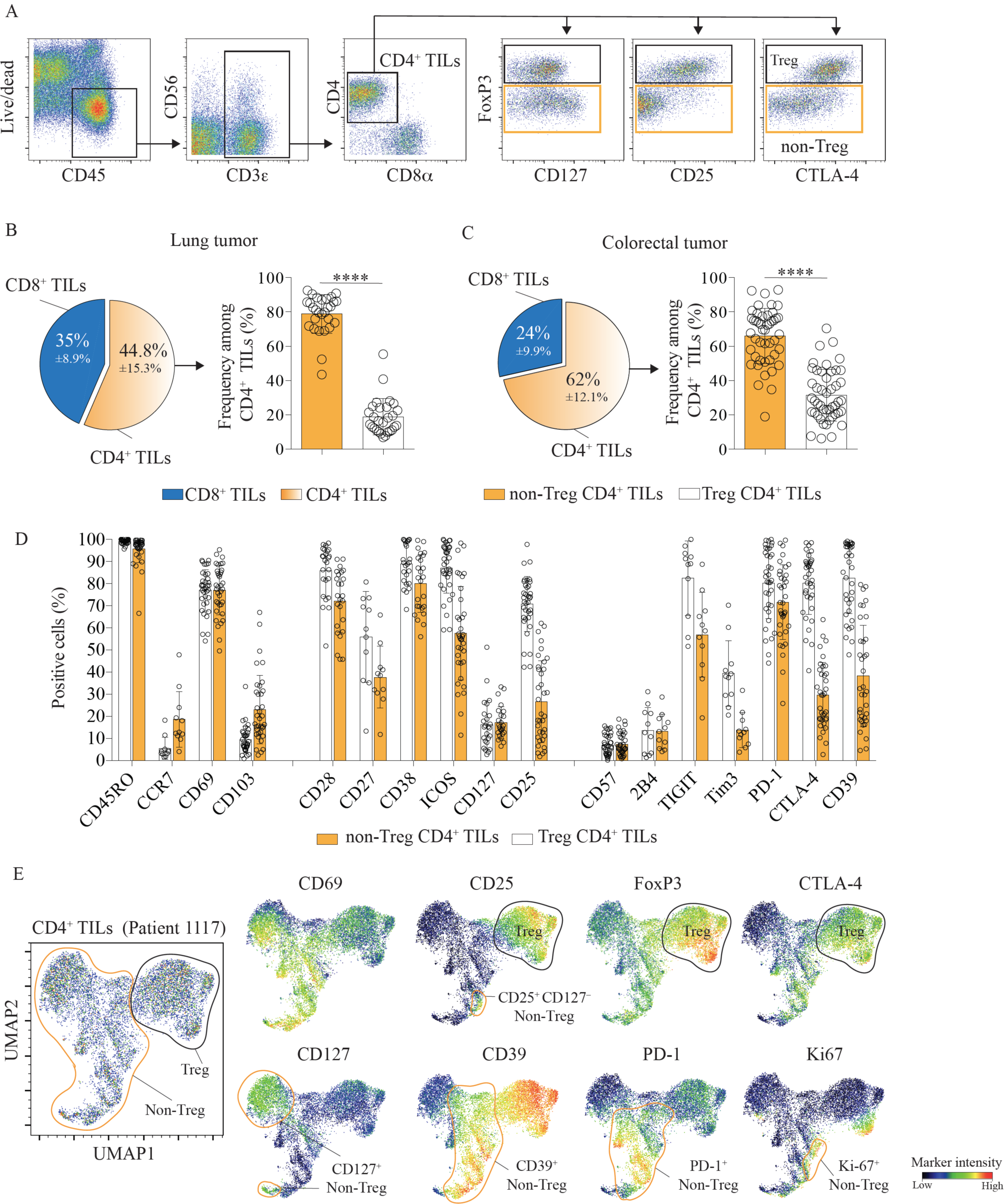
CD4+ TILs are composed of a majority of non-Treg cells with a heterogeneous phenotypic profile. **A**. Gating strategy to distinguish between Treg (Live CD45+ CD3i:,+ CD4+ Foxp3+) and Non-Treg CD4+ TILs (FoxP3-). Representative mass-cytometry data from one colorectal cancer patient. **B**. Frequency of CD4+ and CDS+ TILs among total CD3+ TILs (left panel) and frequency ofNon-Treg CD4+ TILs vs. Treg cells in lung cancer (right panel), n=28 patients. Data from at least 10 independent experiments using mass cytometry. Means± SD. **C**. Frequency of CD4+ and CDS+ TILs among total CD3+ TILs (left panel) and frequency ofNon-Treg CD4+ TILs and Treg cells in colorectal cancer (right panel), n=5 l patients. Data from at least 10 independent experiments using mass cytometry. Means ± SD. **D**. Expression of selected markers by Non-Treg CD4+ (orange) and Treg (white) TILs in colorectal tumors (n = 25-36 biologically independent individuals). Data from at least 10 independent mass cytometry experi ments. Means± SD, Paired t test - two-tailed. **E**. UMAP plot on total CD4+ TILs exported from a representative colorectal tumor sample. UMAP analysis was performed to explore the heterogeneity of non-Treg CD4+ TILs at the individual level (see also Figure S2B).

After exploring the diversity of non-Treg CD4^+^ TILs across patients, we performed UMAP analysis to explore the heterogeneity of CD4^+^ TILs within individuals. In one example, we distinguished several cell clusters, illustrating a broad phenotypic heterogeneity (Figure 2E and S2B). We first identified a cell population with Treg cells features (FoxP3^+^, CD25^+^, CD127^−^, CTLA-4^+^). Among the non-Treg CD4^+^ TILs, we observed presence of multiple cell clusters expressing stimulatory and inhibitory markers at variable intensities. For instance, CD127 (a.k.a IL-7R) that promotes survival of effector cells, could only be found in some of the clusters. Within the cell clusters expressing CD39, we detected differential expression levels of inhibitory receptors such as PD-1, CTLA-4 and Ki-67 suggesting an ongoing antigen exposure and cell expansion (Figure 2E and S3).

Overall, these data showed a high degree of phenotypic diversity among non-Treg CD4^+^ TILs within individual tumors and across patients. Phenotypic analysis showed that both effectors and exhausted cells were found at the same time in the same tumor.

### Cancer-unrelated non-Treg CD4^+^ TILs infiltrate tumor and lack CD39 expression

As we and others have shown that cancer-unrelated bystander CD8^+^ TILs are abundant in cancer and phenotypically distinct (i.e. lack of CD39 expression) (7-9), we explored whether CD39^−^ non-Treg CD4^+^ TILs could be also enriched for cancer unrelated antigen-specific cells. Strikingly, we observed an important heterogeneity for CD39 expression across both cohorts, with patients showing up to 95% of CD39^−^ non-Treg CD4^+^ TILs and others showing less than 20% (Figure 3A, 3B and 3C). We performed a correlation analysis comparing frequencies of CD39^−^ non-Treg CD4^+^ TILs with CD39^−^ CD8^+^ TILs of the same patient (Figure 3B and 3C). In both tumor types, we observed that frequencies of bystander CD8^+^ TILs strongly correlate with the frequency of CD39^−^ non-Treg CD4^+^ TILs. We hypothesized that if CD39^−^ non-Treg CD4^+^ TILs were bystander, they should express a different phenotypic profile. By looking at inhibitory receptors associated with chronic antigen stimulation, we observed a significantly lower expression of TIGIT, CTLA-4 and PD-1 on CD39^−^ non-Treg CD4^+^ TILs as compared to their CD39^+^ counterparts (Figure 3D, 3E and S3). Functionally, CD39^−^ non-Treg CD4^+^ TILs produced more of TNFα and IL-2, suggesting that these cells are more functionally capable and less exhausted (Figure 3F, 3G and S3).

**Figure 3.**
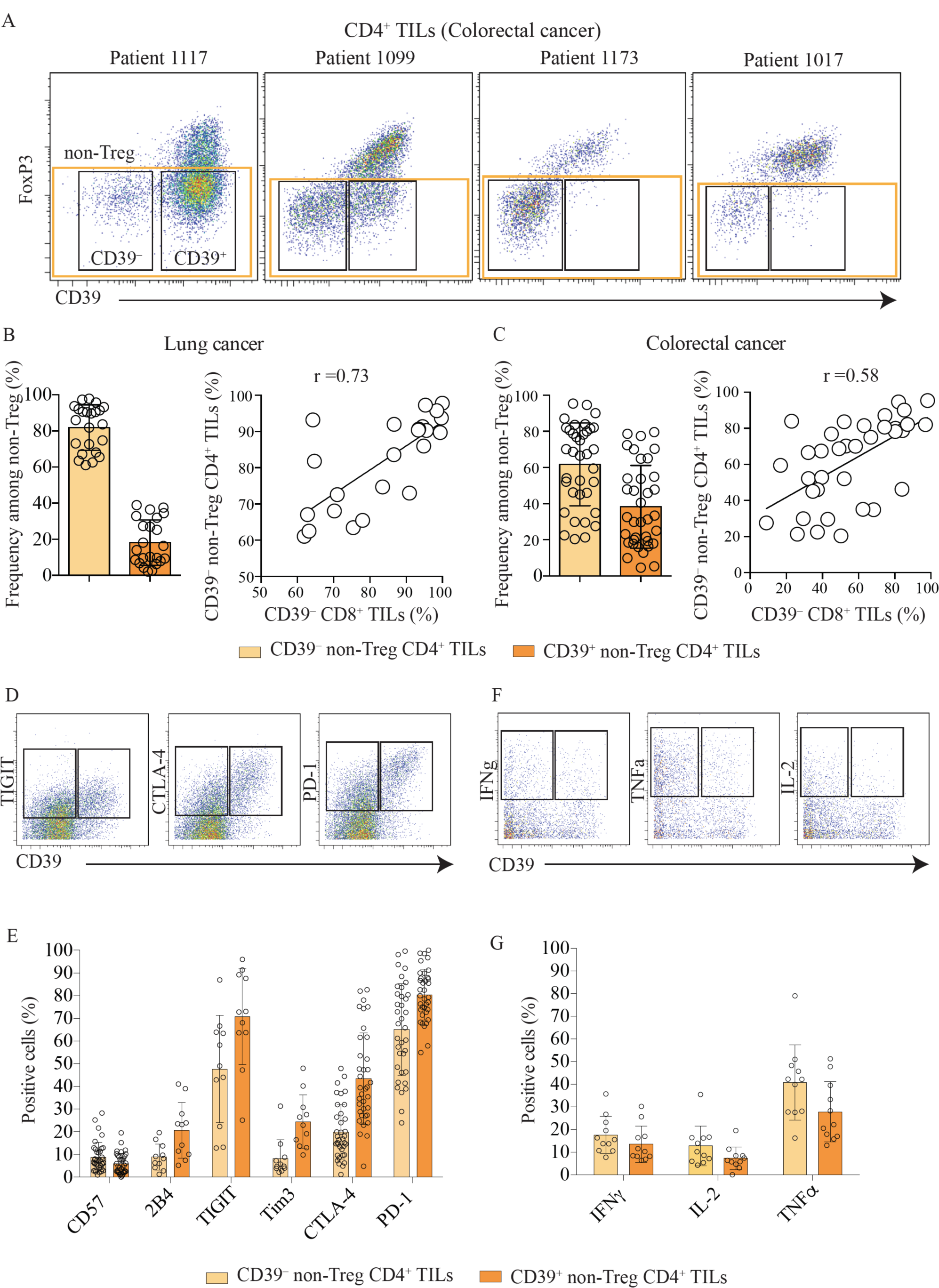
CD39-non-Treg CD4+ TILs are phenotypically and functionally not exhausted. **A**. Dot plots showing differential expression ofCD39 vs. Foxp3 in CD4+ TILs of4 colorectal cancer patients. **B**. CD39 expression on Non-Treg CD4+ vs. Treg TILs (left panel) and correlation between CD39-Non-Treg CD4+ TILs and CD39-CD8+ TILs in lung cancer (right panel), n=21. **C**. CD39 expression on Non-Treg CD4+ vs. Treg TILs (left panel) and correlation between CD39-Non-Treg CD4+ TILs and CD39+ CD8+ TILs in lung cancer (right panel). **D**. Representative staining showing expression ofCD39 vs. inhibitory receptors (TIGIT, CTLA-4 and PD-1) on Non-Treg CD4+TILs in colorectal cancer. **E**. Comparison of inhibitory receptors expression between CD39- and CD39+ Non-Treg CD4+ TILs in colorectal cancer (See Figure S3 for lung cancer). **F**. Representative staining comparing expression ofCD39 vs. cytokine production (IFNγ, TNFα and IL-2) on Non-Treg CD4+TILs in colorectal cancer. **G.** Comparison ofcytokine production between CD39- and CD39+ Non-Treg CD4+ TILs in colorectal cancer (See Figure S3 for lung cancer).

To confirm our hypothesis of bystander CD4^+^ TILs, we first screened tumor tissues with MHC class II tetramers specific for allergen, tumor antigens, EBV or Flu epitopes. Even though we detected these cells in blood after tetramer enrichment (Figure S4), we failed to detect them in tumor tissues (see Discussion). In order to bypass the use of tetramers to assess presence of CD4^+^ T cells specific for cancer unrelated antigens in the tumors, we optimized an activation-induced marker (AIM) assay to assess activation of CD4^+^ TILs stimulated with cancer-unrelated epitopes (here HCMV peptide pool, see methods)(35)(Figure 4A). By measuring the up-regulation of both CD40L and CD69, we observed the presence of HCMV-specific CD4^+^ TILs from the tumors (Figure 4B). When compared with the paired CD4^+^ T cells from PBMC, we observed a higher frequency and fold change of HCMV-specific cells in CD4^+^ TILs, showing that similarly to CD8^+^ TILs, cancer-unrelated CD4^+^ T cells infiltrate tumor tissues (Figure 4C and D). These cells also lacked CD39 expression when analyzed together with total CD4^+^ TILs (Figure 4E), suggesting that the lack of CD39 could also be a straightforward marker to identify non-Treg cancer-unrelated CD4^+^TILs.

**Figure 4.**
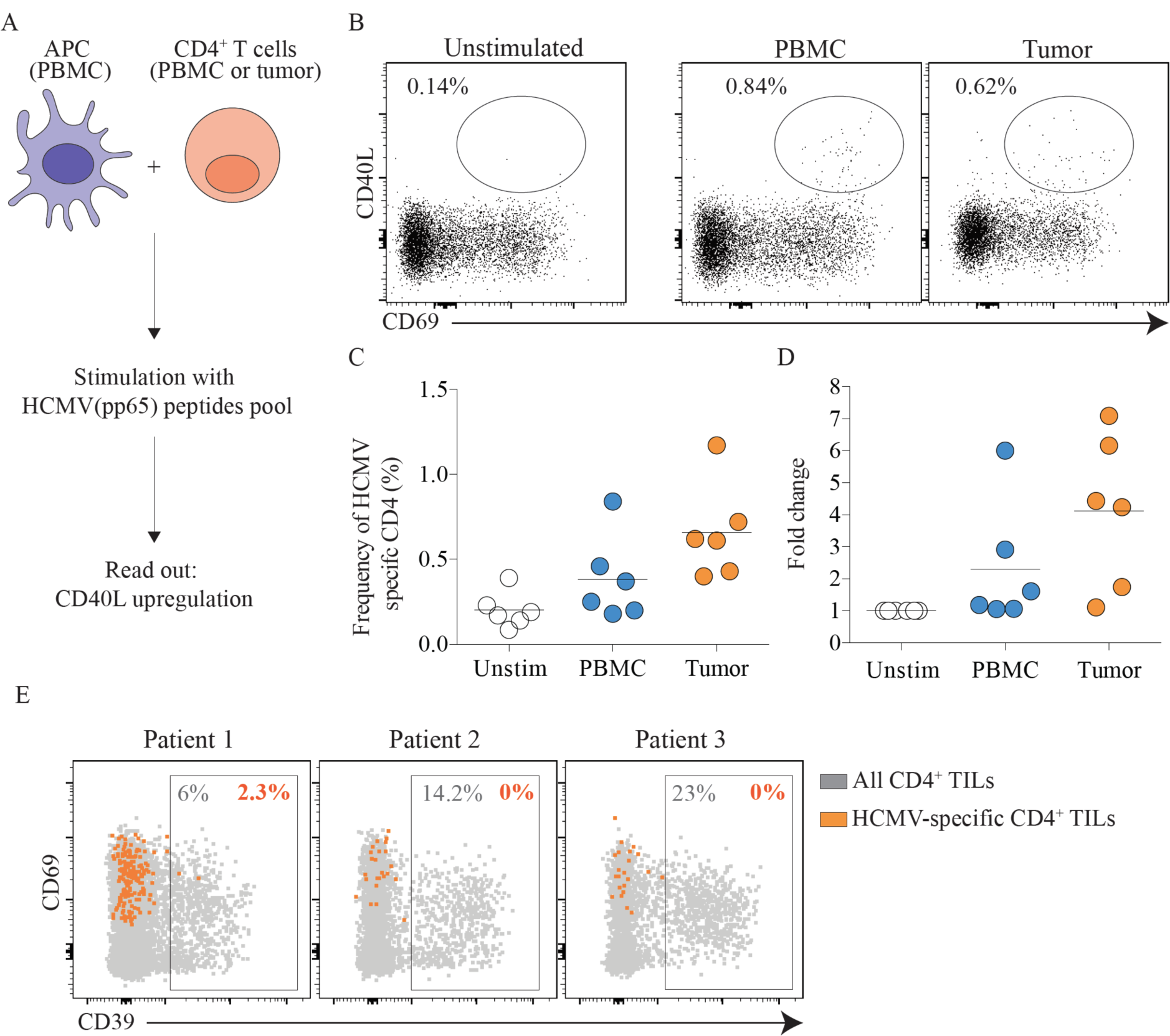
Bystander CD4+ TILs infiltrate tumor and lack expression of CD39. **A**. Schematic for in-vitro activation induced marker (AIM) assay to assess the reactivity of CD4+ TILs toHCMV pp65 epitopes as previously described (see methods, (35). APC and CD4 T cells were isolated from the same donor. **B**. Representative staining showing expression of CD40L and CD69 by CD4+ T cells from PBMC or tumor sample after overnight stimulation with HCMV pp65 peptides pool (see methods). **C**. Frequency of HCMV (pp65)-specific CD4+ T cells in PBMC and tumor. n=6 patients, **D**. Fold change ofHCMV (pp65)-specific CD4+ T cells in PBMC and tumor compared to unstimulated. N=6 patients. **E**. Representative dot plots showing expression of CD39 byHCMV-specific CD4+ TILs (orange) vs. total CD4+ TILs (grey) from three lung cancer patients.

## Discussion

Since the late 1990s, research has highlighted the central role of T cells in antitumor immunity (36). Notably, because of their ability to directly kill tumor cells and a better knowledge of MHC class I tumor antigens, much more attention has been dedicated to the role of CD8^+^ T cells (37-39). In the meantime, many studies have also elucidated the detrimental role of CD4^+^ Treg cells in antitumor immunity and put these cells at the center stage as immunotherapy targets (40). Our work brings to light that consistently large fractions of total T cells infiltrating the tumor are made up of non-Treg CD4^+^ T cells in both colorectal and lung cancer. Similar observation has been previously made in breast cancer (14). In lymph nodes, non-Treg CD4^+^ T cells support the priming of tumor-specific CD8^+^ T cells (41). In tumor microenvironment, these cells enhance the activity of CD8^+^ TILs by producing cytokines (i.e. TNFα, IFNγ) but can also act as effectors by eliminating tumor cells in a direct or indirect way (42, 43). Contrary to MHC class I which is expressed by tumor cells and presents tumor antigens to CD8^+^ TILs, MHC class II is usually not expressed (or expressed at low levels) by human tumor cells (44). However, we clearly observe an up-regulation of markers associated with chronic antigen exposure in non-Treg CD4^+^ TILs, such as Ki-67, PD-1, CTLA-4 indicating that these cells can be activated at the tumor site as well (3). We hypothesize that this activation might be mediated by antigen presenting cells, such as macrophages and dendritic cells. The distinct phenotype of non-Treg CD4^+^ TILs observed across patients, especially regarding expression of inhibitory receptors, could be explained by tumor-intrinsic factors shaping the individual tumor immune microenvironment (45). Furthermore, we also observe heterogeneity of non-Treg CD4^+^ TILs within the same tumor, with cells showing an effector phenotype and others expressing hallmarks of chronic antigen stimulation, notably CD39.

CD39 is an enzyme that converts extracellular ATP to AMP. In turn, CD73 converts AMP into adenosine, shown to possess immunosuppressive activity (46). Conversion of extracellular ATP in adenosine by CD39 thus leads to inhibition of CD4, CD8, NK cell function, decreased phagocytosis and antigens presentation activities by macrophages and dendritic cells (47, 48). Widely reported in Treg-related literature, CD39 has also been described on HIV-, HBV- and tumor-specific CD8^+^ T cells as a marker expressed during chronic antigen stimulation (7, 49-51). Yet, only few groups have characterized this marker on non-Treg CD4^+^ TILs. *In-vitro*, CD39 is expressed on Non-Treg CD4^+^ TILs after activation and on Listeria-specific CD4^+^ T cells after infection (32). Interestingly, a pioneer study reported an increased frequency of pathogenic CD39^+^ non-Treg CD4^+^ T cells in the peripheral blood of patients with renal allograft rejection (52). As previously observed for CD8^+^ TILs, CD39 could be a useful marker to identify tumor-specific CD4^+^ T cells as well within the tumor micro-environment. Additional studies will be needed to confirm this hypothesis and to better understand the regulation of CD39 in non-Treg CD4^+^ TILs.

By investigating the antigen specificity of CD4^+^ TILs, we failed to detect MHC class II tetramer positive cells in the tumors. This negative result could be attributed to the limited number of tetramers tested, the low frequency of specific T cells for any given epitope combined with the low number of cells obtained from tumor dissociation(53). Using the AIM assay, we detected cancer unrelated CD4^+^ TILs. These HCMV-specific cells lack CD39 expression, which mirrors our previous observations with CD39^−^ CD8^+^ TILs specific for cancer unrelated antigens (HCMV, EBV, Flu) (7). Of note, the observation that tumor-specific CD4 and CD8 responses are coordinated is consistent with the notion that tumor-specific CD4 responses are also required for the induction of tumor-specific CD8 response as recently illustrated in mice (22). Besides, up to 95% of non-Treg CD4^+^ TILs lack CD39 expression in some patients. Taken together, these two observations could suggest that the majority of effectors TILs are not tumor-specific. This hypothesis could explain, along with other factors, the absence of response in most patients treated with anti-PD-1 (54). Bystander CD4^+^ (and CD8^+^) TILs are in fact not passive in the tumor microenvironment, and several reports have highlighted their role in modulating disease severity upon TCR-independent activation (55, 56). Because of their TCR specificity for known viral epitopes, virus-specific bystander TILs could also be specifically targeted by therapeutic approaches to produce cytokines and enhance anti-tumor response (11). Overall, our findings highlight that non-Treg CD4^+^ TILs cells represent one of the main lymphocytes recruited at the tumor site and as well a potential target of interest for immunotherapy.

## Methods

### Human samples

PBMC and tumor samples were obtained from patients with colorectal cancer or lung cancer. The use of human tissues was approved by the appropriate institutional research boards, A*STAR and the Singapore Immunology Network, Singapore.

### Cell isolation

Samples were prepared as previously described (57). In brief, tissues were mechanically dissociated into small pieces and incubated at 37 °C for 15 to 40 min in DMEM + collagenase IV (1 mg/ml) + DNase (15 μg/ml). Digestion was stopped by addition of RPMI containing 5% FBS. Dissociated tissues were filtered and washed in RPMI 5% + DNase (15 μg/ml) FBS. All samples were cryopreserved in 90% FBS + 10% DMSO and stored in liquid nitrogen.

### Single-cell Sequencing

Experiment was performed as previously described (29). In brief, frozen samples were thawed and washed in RPMI 10% FBS + DNAse (15 ug/ml). Samples were depleted of tumor cells (αEpCAM – clone 9C4), Myeloid cells (αCD14 – clone TUK4) and B cells (αCD20 – clone 2H7) using anti-Mouse IgG microbeads (Miltenyi – 130-048-401). Cells were then incubated with BD AbSeq Ab-oligos following manufacturers’ instructions. Single cells were isolated using Single Cell Capture and cDNA synthesis with the BD Rhapsody Express Single-cell Analysis System. Parallel RNA and BD AbSeq sequencing libraries were generated using BD Rhapsody targeted mRNA (BD – 633751) and AbSeq amplification and BD Single-cell Multiplexing kits and protocol (BD – 633771). Quality of final libraries was assessed using Agilent 2200 TapeStation with High Sensitivity D5000 ScreenTape, quantified using a Qubit Fluorometer (ThermoFisher), and carried through to sequencing with Novaseq S1 on Illumina sequencer. FASTQ files containing sequenced data were analyzed using the Seven Bridges platform provided by BD (See “BD Single Cell Genomics Bioinformatics Handbook – 54169 Rev. 6.0” for specific details) (29).

### Mass-cytometry staining

Samples were stained as previously described (57, 58). In brief, antibody conjugation was performed according to the protocol provided by Fluidigm (See Table S1 for clone list and metals). Prior to surface staining, cells were stained with Cisplatin (viability marker) 5 μM in PBS for 5 min. Cells were then stained in PBS + 0.5% BSA buffer with surface antibodies at 4°C for 15 min. After two washing steps, cells were fixed in fixation FoxP3 buffer (eBioscience – 00-5521-00) for 30 min at 4°C. After washing in perm buffer cells were stained with biotinylated FoxP3 during 30 min at 4°C in perm buffer. Cells were washed and stained with streptavidin coupled to heavy metal for 30mn at 4°C in perm buffer. After two washing steps, cells were fixed in PBS 2% PFA overnight. Prior to CyTOF acquisition, cells were stained for DNA (Cell-ID intercalator-Ir, Fluidigm) for 10 min at room temperature, washed three times with dH20 and acquired on CyTOF.

### Data analysis and UMAP

After mass cytometry (CyTOF) acquisition, any zero values were randomized using a uniform distribution of values between 0 and −1 using R. The signal of each parameter was normalized based on EQ beads (Fluidigm) as described previously (59). Samples were then used for UMAP analysis similar to that previously described using customized R scripts based on the ‘flowCore’ and ‘uwot’ R packages (31). In R, all data were transformed using the logicleTransform function (flowCore package) using parameters: *w* = 0.25, *t* = 16409, *m* = 4.5, *a* = 0 to roughly match scaling historically used in FlowJo. For heatmaps, median intensity corresponds to a logical data scale using formula previously described (60). The colors in the heat map represent the measured means intensity value of a given marker in a given sample. A seven-color scale is used with black–blue indicating low expression values, green–yellow indicating intermediately expressed markers, and orange-red representing highly expressed markers. Violin plots were generated using customized R scripts based on the ‘ggplot2’ R package (geom_violin, geom_boxplot, geom_quasirandom).

### AIM (activation induced marker) assay

AIM assay was performed as described previously (35). Briefly, on day 1, frozen paired blood and tumor samples were thawed and prepared as stated above. APC (gated as all CD3-CD45^+^live) were sorted from the PBMC, CD4^+^ T cells (gated as CD45^+^live CD3^+^CD4^+^) were sorted from blood and the tumors using BD FACSAria II. After sorting, cells were rested for 3h at 37°C, incubated with a CD40 blocking antibody for 15 min and put in coculture at a ratio of 1 CD4: 5 APC. Cells were then stimulated with either HCMV peptides pool (Catalogue number, 86.25ug/ml), DMSO (negative control, 100ug/ml) or SEB (positive control, 500ug/ml) for 18h. On day 2, cells were washed, stained with surface flow antibodies (Table S2) and acquired on BD FACSCelesta. Activation was measured with CD69 and CD40L expression on total CD4^+^ T cells and bystander CD4^+^ T cells were analyzed for CD39 expression.

## References

1. Dunn GP, Bruce AT, Ikeda H, Old LJ, Schreiber RD. Cancer immunoediting: from immunosurveillance to tumor escape. Nat Immunol. 2002;3(11):991–8.

2. Schreiber RD, Old LJ, Smyth MJ. Cancer immunoediting: integrating immunity’s roles in cancer suppression and promotion. Science. 2011;331(6024):1565–70.

3. Pardoll DM. The blockade of immune checkpoints in cancer immunotherapy. Nat Rev Cancer. 2012;12(4):252–64.

4. Rosenberg SA, Restifo NP. Adoptive cell transfer as personalized immunotherapy for human cancer. Science. 2015;348(6230):62–8.

5. Ott PA, Hu Z, Keskin DB, Shukla SA, Sun J, Bozym DJ, et al. An immunogenic personal neoantigen vaccine for patients with melanoma. Nature. 2017;547(7662):217–21.

6. Maleki Vareki S, Garrigos C, Duran I. Biomarkers of response to PD-1/PD-L1 inhibition. Crit Rev Oncol Hematol. 2017;116:116–24.

7. Simoni Y, Becht E, Fehlings M, Loh CY, Koo SL, Teng KWW, et al. Bystander CD8(+) T cells are abundant and phenotypically distinct in human tumour infiltrates. Nature. 2018;557(7706):575–9.

8. Canale FP, Ramello MC, Nunez N, Araujo Furlan CL, Bossio SN, Gorosito Serran M, et al. CD39 Expression Defines Cell Exhaustion in Tumor-Infiltrating CD8(+) T Cells. Cancer Res. 2018;78(1):115–28.

9. Duhen T, Duhen R, Montler R, Moses J, Moudgil T, de Miranda NF, et al. Co-expression of CD39 and CD103 identifies tumor-reactive CD8 T cells in human solid tumors. Nat Commun. 2018;9(1):2724.

10. Scheper W, Kelderman S, Fanchi LF, Linnemann C, Bendle G, de Rooij MAJ, et al. Low and variable tumor reactivity of the intratumoral TCR repertoire in human cancers. Nat Med. 2019;25(1):89–94.

11. Rosato PC, Wijeyesinghe S, Stolley JM, Nelson CE, Davis RL, Manlove LS, et al. Virus-specific memory T cells populate tumors and can be repurposed for tumor immunotherapy. Nat Commun. 2019;10(1):567.

12. Wing JB, Tanaka A, Sakaguchi S. Human FOXP3(+) Regulatory T Cell Heterogeneity and Function in Autoimmunity and Cancer. Immunity. 2019;50(2):302–16.

13. Saito T, Nishikawa H, Wada H, Nagano Y, Sugiyama D, Atarashi K, et al. Two FOXP3(+)CD4(+) T cell subpopulations distinctly control the prognosis of colorectal cancers. Nat Med. 2016;22(6):679–84.

14. Plitas G, Konopacki C, Wu K, Bos PD, Morrow M, Putintseva EV, et al. Regulatory T Cells Exhibit Distinct Features in Human Breast Cancer. Immunity. 2016;45(5):1122–34.

15. Facciabene A, Motz GT, Coukos G. T-regulatory cells: key players in tumor immune escape and angiogenesis. Cancer Res. 2012;72(9):2162–71.

16. Zou W. Regulatory T cells, tumour immunity and immunotherapy. Nat Rev Immunol. 2006;6(4):295–307.

17. Tang F, Du X, Liu M, Zheng P, Liu Y. Anti-CTLA-4 antibodies in cancer immunotherapy: selective depletion of intratumoral regulatory T cells or checkpoint blockade? Cell Biosci. 2018;8:30.

18. Callahan MK, Wolchok JD, Allison JP. Anti-CTLA-4 antibody therapy: immune monitoring during clinical development of a novel immunotherapy. Semin Oncol. 2010;37(5):473–84.

19. Seidel JA, Otsuka A, Kabashima K. Anti-PD-1 and Anti-CTLA-4 Therapies in Cancer: Mechanisms of Action, Efficacy, and Limitations. Front Oncol. 2018;8:86.

20. Arce Vargas F, Furness AJS, Solomon I, Joshi K, Mekkaoui L, Lesko MH, et al. Fc-Optimized Anti-CD25 Depletes Tumor-Infiltrating Regulatory T Cells and Synergizes with PD-1 Blockade to Eradicate Established Tumors. Immunity. 2017;46(4):577–86.

21. Arce Vargas F, Furness AJS, Litchfield K, Joshi K, Rosenthal R, Ghorani E, et al. Fc Effector Function Contributes to the Activity of Human Anti-CTLA-4 Antibodies. Cancer Cell. 2018;33(4):649–63 e4.

22. Alspach E, Lussier DM, Miceli AP, Kizhvatov I, DuPage M, Luoma AM, et al. MHC-II neoantigens shape tumour immunity and response to immunotherapy. Nature. 2019;574(7780):696–701.

23. Melssen M, Slingluff CL, Jr. Vaccines targeting helper T cells for cancer immunotherapy. Curr Opin Immunol. 2017;47:85–92.

24. Quezada SA, Simpson TR, Peggs KS, Merghoub T, Vider J, Fan X, et al. Tumor-reactive CD4(+) T cells develop cytotoxic activity and eradicate large established melanoma after transfer into lymphopenic hosts. J Exp Med. 2010;207(3):637–50.

25. Wong SB, Bos R, Sherman LA. Tumor-specific CD4+ T cells render the tumor environment permissive for infiltration by low-avidity CD8+ T cells. J Immunol. 2008;180(5):3122–31.

26. Church SE, Jensen SM, Antony PA, Restifo NP, Fox BA. Tumor-specific CD4+ T cells maintain effector and memory tumor-specific CD8+ T cells. Eur J Immunol. 2014;44(1):69–79.

27. Matsuzaki J, Tsuji T, Luescher IF, Shiku H, Mineno J, Okamoto S, et al. Direct tumor recognition by a human CD4(+) T-cell subset potently mediates tumor growth inhibition and orchestrates anti-tumor immune responses. Sci Rep. 2015;5:14896.

28. Malandro N, Budhu S, Kuhn NF, Liu C, Murphy JT, Cortez C, et al. Clonal Abundance of Tumor-Specific CD4(+) T Cells Potentiates Efficacy and Alters Susceptibility to Exhaustion. Immunity. 2016;44(1):179–93.

29. Mair F, Erickson JR, Voillet V, Simoni Y, Bi T, Tyznik AJ, et al. A Targeted Multi-omic Analysis Approach Measures Protein Expression and Low-Abundance Transcripts on the Single-Cell Level. Cell Rep. 2020;31(1):107499.

30. Shahi P, Kim SC, Haliburton JR, Gartner ZJ, Abate AR. Abseq: Ultrahigh-throughput single cell protein profiling with droplet microfluidic barcoding. Sci Rep. 2017;7:44447.

31. Becht E, McInnes L, Healy J, Dutertre CA, Kwok IWH, Ng LG, et al. Dimensionality reduction for visualizing single-cell data using UMAP. Nat Biotechnol. 2018.

32. Raczkowski F, Rissiek A, Ricklefs I, Heiss K, Schumacher V, Wundenberg K, et al. CD39 is upregulated during activation of mouse and human T cells and attenuates the immune response to Listeria monocytogenes. PLoS One. 2018;13(5):e0197151.

33. Yan D, Farache J, Mingueneau M, Mathis D, Benoist C. Imbalanced signal transduction in regulatory T cells expressing the transcription factor FoxP3. Proc Natl Acad Sci U S A. 2015;112(48):14942–7.

34. Han S, Yang Y. Interleukin-32: Frenemy in cancer? BMB Rep. 2019;52(3):165–74.

35. Reiss S, Baxter AE, Cirelli KM, Dan JM, Morou A, Daigneault A, et al. Comparative analysis of activation induced marker (AIM) assays for sensitive identification of antigen-specific CD4 T cells. PLoS One. 2017;12(10):e0186998.

36. Shankaran V, Ikeda H, Bruce AT, White JM, Swanson PE, Old LJ, et al. IFNgamma and lymphocytes prevent primary tumour development and shape tumour immunogenicity. Nature. 2001;410(6832):1107–11.

37. Reading JL, Galvez-Cancino F, Swanton C, Lladser A, Peggs KS, Quezada SA. The function and dysfunction of memory CD8(+) T cells in tumor immunity. Immunol Rev. 2018;283(1):194–212.

38. Farhood B, Najafi M, Mortezaee K. CD8(+) cytotoxic T lymphocytes in cancer immunotherapy: A review. J Cell Physiol. 2019;234(6):8509–21.

39. van der Leun AM, Thommen DS, Schumacher TN. CD8(+) T cell states in human cancer: insights from single-cell analysis. Nat Rev Cancer. 2020;20(4):218–32.

40. Tanaka A, Sakaguchi S. Targeting Treg cells in cancer immunotherapy. Eur J Immunol. 2019;49(8):1140–6.

41. Ahrends T, Spanjaard A, Pilzecker B, Babala N, Bovens A, Xiao Y, et al. CD4(+) T Cell Help Confers a Cytotoxic T Cell Effector Program Including Coinhibitory Receptor Downregulation and Increased Tissue Invasiveness. Immunity. 2017;47(5):848–61 e5.

42. Perez-Diez A, Joncker NT, Choi K, Chan WF, Anderson CC, Lantz O, et al. CD4 cells can be more efficient at tumor rejection than CD8 cells. Blood. 2007;109(12):5346–54.

43. Mumberg D, Monach PA, Wanderling S, Philip M, Toledano AY, Schreiber RD, et al. CD4(+) T cells eliminate MHC class II-negative cancer cells in vivo by indirect effects of IFN-gamma. Proc Natl Acad Sci U S A. 1999;96(15):8633–8.

44. Haabeth OA, Tveita AA, Fauskanger M, Schjesvold F, Lorvik KB, Hofgaard PO, et al. How Do CD4(+) T Cells Detect and Eliminate Tumor Cells That Either Lack or Express MHC Class II Molecules? Front Immunol. 2014;5:174.

45. Li J, Byrne KT, Yan F, Yamazoe T, Chen Z, Baslan T, et al. Tumor Cell-Intrinsic Factors Underlie Heterogeneity of Immune Cell Infiltration and Response to Immunotherapy. Immunity. 2018;49(1):178–93 e7.

46. Bastid J, Cottalorda-Regairaz A, Alberici G, Bonnefoy N, Eliaou JF, Bensussan A. ENTPD1/CD39 is a promising therapeutic target in oncology. Oncogene. 2013;32(14):1743–51.

47. Elliott MR, Chekeni FB, Trampont PC, Lazarowski ER, Kadl A, Walk SF, et al. Nucleotides released by apoptotic cells act as a find-me signal to promote phagocytic clearance. Nature. 2009;461(7261):282–6.

48. Ohta A, Gorelik E, Prasad SJ, Ronchese F, Lukashev D, Wong MK, et al. A2A adenosine receptor protects tumors from antitumor T cells. Proc Natl Acad Sci U S A. 2006;103(35):13132–7.

49. Gupta PK, Godec J, Wolski D, Adland E, Yates K, Pauken KE, et al. CD39 Expression Identifies Terminally Exhausted CD8+ T Cells. PLoS Pathog. 2015;11(10):e1005177.

50. Deaglio S, Dwyer KM, Gao W, Friedman D, Usheva A, Erat A, et al. Adenosine generation catalyzed by CD39 and CD73 expressed on regulatory T cells mediates immune suppression. J Exp Med. 2007;204(6):1257–65.

51. Borsellino G, Kleinewietfeld M, Di Mitri D, Sternjak A, Diamantini A, Giometto R, et al. Expression of ectonucleotidase CD39 by Foxp3+ Treg cells: hydrolysis of extracellular ATP and immune suppression. Blood. 2007;110(4):1225–32.

52. Dwyer KM, Hanidziar D, Putheti P, Hill PA, Pommey S, McRae JL, et al. Expression of CD39 by human peripheral blood CD4+ CD25+ T cells denotes a regulatory memory phenotype. Am J Transplant. 2010;10(11):2410–20.

53. Nepom GT. MHC class II tetramers. J Immunol. 2012;188(6):2477–82.

54. Chen DS, Mellman I. Elements of cancer immunity and the cancer-immune set point. Nature. 2017;541(7637):321–30.

55. Whiteside SK, Snook JP, Williams MA, Weis JJ. Bystander T Cells: A Balancing Act of Friends and Foes. Trends Immunol. 2018;39(12):1021–35.

56. Christoffersson G, Chodaczek G, Ratliff SS, Coppieters K, von Herrath MG. Suppression of diabetes by accumulation of non-islet-specific CD8(+) effector T cells in pancreatic islets. Sci Immunol. 2018;3(21).

57. Simoni Y, Fehlings M, Kloverpris HN, McGovern N, Koo SL, Loh CY, et al. Human Innate Lymphoid Cell Subsets Possess Tissue-Type Based Heterogeneity in Phenotype and Frequency. Immunity. 2017;46(1):148–61.

58. Simoni Y, Fehlings M, Newell EW. Multiplex MHC Class I Tetramer Combined with Intranuclear Staining by Mass Cytometry. Methods Mol Biol. 2019;1989:147–58.

59. Finck R, Simonds EF, Jager A, Krishnaswamy S, Sachs K, Fantl W, et al. Normalization of mass cytometry data with bead standards. Cytometry A. 2013;83(5):483–94.

60. Moore WA, Parks DR. Update for the logicle data scale including operational code implementations. Cytometry A. 2012;81(4):273–7.

